# Insect metamorphosis is regulated differently between sexes by members of a microRNA cluster

**DOI:** 10.1101/2024.06.25.600558

**Authors:** Chade Li, Ki Kei Chan, Wenyan Nong, ShanShan Chen, Wai Lok So, Zhe Qu, Heidi Y.C. Wu, Ho Yin Yip, Chi Bun Chan, Stephen S. Tobe, William G. Bendena, Zhen Peng Kai, Jerome H.L. Hui

## Abstract

Insects comprise the majority of all described animal species and dominate the terrestrial habitats. The evolution of insect metamorphosis played a profound role in their successful adaptation and radiation. Insect metamorphosis is dependent on hormones ecdysteroids and sesquiterpenoids such as juvenile hormone. Despite the fact there are genuine differences between sexes during insect metamorphosis which facilitate their successful mating, how such sexual dimorphism in metamorphosis is being controlled is poorly known. We first generated transcriptomic profiles of male and female flies in late larvae and early pupae stages. Using a combination of genome-wide prediction and *in vitro* dual-luciferase validations, members of a microRNA cluster miR-277/34 were found to potentially regulate the neuropeptide receptor (*AstC-R1*) that when activated inhibits the sesquiterpenoid pathway and a juvenile hormone-dependent transcription factor (*Kr-h1*) in fly *Drosophila melanogaster.* Loss-of-function mutants were created deleting either miR-277 or miR-34, and expression levels of both *AstC-R1* and *Kr-h1* as well as ecdysteroid and sesquiterpenoid hormone titres were altered. Further comparison of transcriptomes of the late larvae and early pupae of both sexes revealed differential gene pathways being regulated by members of miR-277/34 between sexes during metamorphosis. This study highlights how members of a microRNA cluster control hormonal and developmental gene pathways in different sexes of insects during metamorphosis.

## Background

The Arthropoda is the largest animal phylum, containing most of the described living species in the world, while the Insecta (hexapods) is the most abundant class of arthropods. Majority of insects undergo metamorphosis, which results in an alteration to the whole body-plan during development (Truman and Riddiford 1999; Belles 2011). Such strategy of changing in body form undeniably played an important role during the evolution and successful radiation of insects, such as reduce competition of food source and increase exploration and settlement in new environment (Cheong et al 2015; Truman 2019).

Insect metamorphosis is strictly programmed and precisely controlled by the hormonal systems. The three major groups of hormones govern metamorphosis include ecdysteroids, neuropeptides, and sesquiterpenoids (Bendena et al 1999; Tobe and Bendena 1999; Truman and Riddiford 2002; Riddiford 2012; Cao et al 2017; Qu et al 2018; Belles 2017; Tsang et al 2020). In general, these three hormones work in a cooperative manner, where ecdysteroid 20-hydroxyecdysone control the timing of ecdysis or molting, neuropeptide ecdysis-triggering hormone act as regulators of ecdysteroid synthesis or release, and sesquiterpenoids such as juvenile hormone (JH) regulate metamorphosis based on their titre levels.

During larva to larva ecdysis, JH binds to intracellular receptor *Methoprene-tolerant* (Met) and subsequently the transcription factor *Krüppel homolog 1* (Kr-h1) to suppress metamorphosis to pupa (Konopova and Jindra 2007; Parthasarathy et al 2008; Minakuchi et al 2009); and during larva to adult metamorphosis, in the absence of JH, transcription factor *E93* induces metamorphosis via the MEKRE93 (Met-Kr-h1-E93) pathway (Ureña et al 2016). MicroRNAs are conserved noncoding RNAs that play crucial roles in many biological processes in insects by post-transcriptional regulation. In the last decade, a number of microRNAs have been identified that contribute to the regulation of insect metamorphosis (e.g. Caygill and Johnston 2008; Sokol et al 2008; Lozano et al 2015).

In the fruit fly *Drosophila melanogaste*r, microRNAs have been found to regulate hormonal and metabolic pathways (e.g. Chan et al 2023; Qu et al 2017). Similar to other holometabolous insects, the life cycle of *Drosophila* exhibits embryo, larva, pupa, and adult stages. Notably, *Drosophila* also shows sexual dimorphism in many aspects, including reproductive behaviours (Han et al 2022), abdomen segments (Wang et al 2011), and responsiveness to ethanol (Devineni and Heberlein 2012). Interestingly, feeding males with juvenile hormone inhibitor could lead to feminization, and this effect is reversible by the simultaneous application juvenile hormone analogue, suggesting hormonal control of sexual dimorphism (Belgacem and Martin 2002).

Furthermore, previous study revealed sex-biased expression of microRNAs in various tissues of males and females flies (Fagegaltier et al 2014). For instance, let-7 expressed in the heads of both sexes, while its expression in the gonads could only be found in the males. Interestingly, let-7 expression could be induced by ecdysone, while the maintenance of sexual identity in male adulthood is regulated by let-7 and ecdysone signalling pathway genes (Sempere et al 2002). However, whether microRNA could be involved in hormonal control and sexual dimorphism during insect metamorphosis, remains unknown.

Here we identified a microRNA cluster involved in metamorphosis by comprehensive mRNA and small RNA sequencing together with genome-wide *in silico* and *in vitro* analyses. Subsequent knockout of members of this microRNA cluster and analyses demonstrated that developmental and hormonal pathway genes were differentially regulated contributing to the sexual dimorphism of insect metamorphosis.

## Results

### Differential protein-coding genes and microRNAs expression in male and female flies during metamorphosis

To reveal the protein-coding genes and microRNAs expression in different sexes of flies during metamorphosis, we have first sequenced the transcriptomes of L3 larvae instars and white pre-pupae from both sexes. A total of 2,548 protein-coding genes were found to be differentially regulated in females during metamorphosis (981 upregulated and 1,567 downregulated), while 2,650 protein-coding genes were differentially regulated in males (1,002 upregulated and 1,648 downregulated) (Figure 1A; Supplementary Table S1, S2; Supplementary Figure S1; Supplementary File 1). We then tested if any functional gene ontology terms were enriched in either sex or during metamorphosis for these differentially expressed genes, and the results are summarised in Supplementary Table S3. In sum, most of the enriched gene pathways are revealed for the downregulated genes during metamorphosis, including the *insect hormone biosynthesis* pathway in both males and females, while some sex-specific enriched gene pathways, such as amino acid metabolic pathways in males and *ascorbate and aldarate metabolism* in females, could also be identified (Figure 1A).

**Figure 1.**
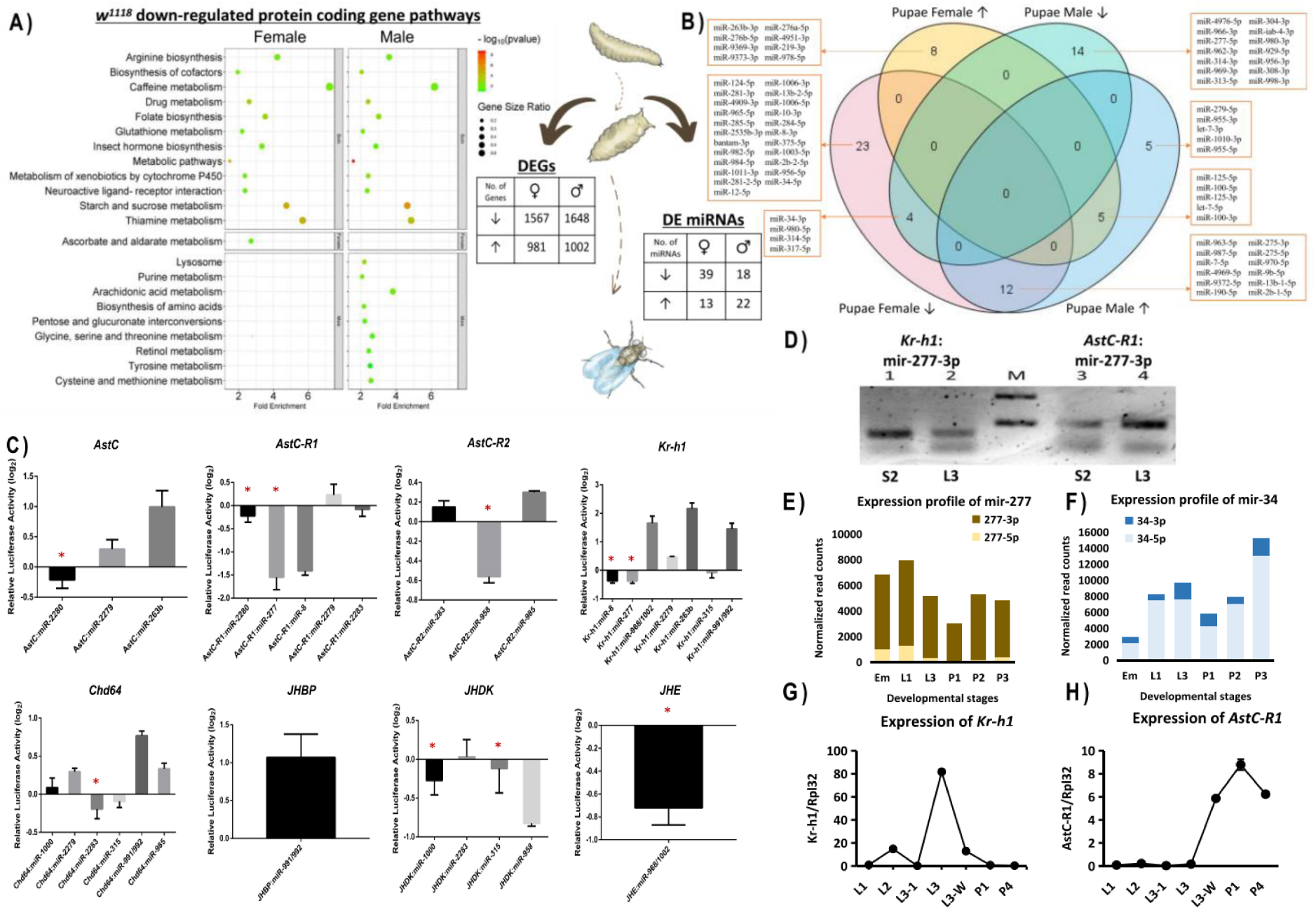
A) Enriched gene pathways and differentially expressed genes (DEGs) of *w^1118^* male and female flies during larvae-pupae transition; B) Differentially expressed microRNAs of *w^1118^* male and female flies in the larvae-pupae transition; C) Screening of potential microRNA candidates that can regulate sesquiterpenoid pathway gene candidates picked up by genome-wide *in silico* prediction; D) MicroRNA pull-down assay showing interactions of miR-277-3p and *Kr-h1* and *AstC-R1* transcripts in *Drosophila* S2 cells and 3^rd^ instar larvae *in vivo*; E-F) Expression profiles of miR-277-3p, miR-277-5p, miR-34-3p and miR-34-5p read counts in fly embryos, larval and pupal stages; G-H) Expression level of *Kr-h1* and *AstC-R1* at larval and pupal stages and pupae stages.

On the other hand, we have also sequenced and uncovered a total of 52 microRNAs was differentially regulated in females during metamorphosis (13 upregulated and 39 downregulated), and 40 microRNAs were differentially regulated in males (22 upregulated and 18 downregulated) (Figure 1B; Supplementary Table S4, S5; Supplementary Figure 2; Supplementary File 1). There were 9 microRNAs that were found to be differentially expressed similarly between sexes during metamorphosis, including miR-314-5p, miR-317-5p, miR-34-3p and miR-980-5p to be downregulated, and miR-100-5p, miR-100-3p, miR-125-5p, miR-125-3p and let-7-5p to be upregulated in both sexes during metamorphosis. In addition, there were 12 microRNAs that showed opposite trends in expression between sexes during metamorphosis (e.g. up-regulated in male but down-regulated in female), including miR-2b-1-5p, miR-7-5p, miR-9b-5p, miR-13b-1-5p, miR-190-5p, miR-275-5p, miR-275-3p, miR-4969-5p, miR-9372-5p, miR-963-5p, miR-970-5p and miR-987-5p (Figure 1B; Supplementary Figure 3 and 4).

### Screening for microRNAs targeting the sesquiterpenoid signalling pathway genes

In insects, sesquiterpenoids is one of the key hormones that regulate metamorphosis, and there are less than a handful of published studies on microRNA regulation of sesquiterpenoids in fly *D. melanogaster*, such as microRNA *bantam* regulates gene transcript of the JHAMT gene and sesquiterpenoid biosynthesis (Qu et al 2017). Using the newly generated transcriptomic data, we further tested if there were other potential microRNAs involved in regulation of other parts of the sesquiterpenoid system. Genome-wide *in silico* prediction of microRNAs that can potentially regulate different genes involved in the sesquiterpenoid system of fly was carried out (Marco et al 2010; Griffiths-Jones et al 2011; Qu et al 2020), and a list of 17 common microRNAs that can potentially regulate the sesquiterpenoid system (other than the biosynthetic pathway) were predicted (Supplementary Figure 5; Supplementary File 1).

Utilising an established cell culture dual-luciferase reporter assay (Hui et al 2013); we have validated 7 microRNAs (miR-8, miR-277, miR-958, miR-1000, miR-1002, miR-2283, miR-2280) that can downregulate the expression of *Drosophila* sesquiterpenoid pathway genes to varying degrees, including *allatostatin C* (*AstC*), *allatostatin C receptor 1* (*AstC-R1*), *allatostatin C receptor 2* (*AstC-R2*), *Krüppel homolog 1* (*Kr-h1*), *Chd64*, *juvenile hormone binding protein (JHBP)*, *juvenile hormone diol kinase* (*JHDK*) and *juvenile hormone esterase* (*JHE*) (Figure 1C). This *in vitro* screening confirms a list of potential microRNAs and sesquiterpenoid gene interactions.

Among these genes, Kr-h1 or Kruppel homolog 1, is a juvenile hormone dependent transcription factor, and have been found to be regulated by another microRNA in cockroaches (Lozano et al 2015); while AstC-R1 or allatostatin C receptor 1, is a G-protein coupled receptor for neuropeptide allatostatin C which regulates juvenile hormone (Stay and Tobe 2007). Interestingly, it can be found that *Kr-h1* and two *AstC* receptors were targets of either or both members of miR-277/34 cluster in our *in vitro* analyses in flies above. To delineate whether miR-277 can selectively bind to AstC-R1 and Kr-h1 transcript in the presence of multiple transcripts *in vivo*, a labelled microRNA pull-down (LAMP) assay system (Hsu et al 2009; Qu et al 2017) was applied, and miR-277 has been shown to directly bind and regulate AstC-R1 and Kr-h1 transcripts *in vivo*, in agreement with the above *in silico* and *in vitro* results (Figure 1D). We also investigated the expression of miR-277 and miR-34 during *D. melanogaster* development with small RNA sequencing and qPCR, which revealed a correlative expression pattern with the expression level of Kr-h1 and AstC-R1 (Figure 1E-H).

### Functional analyses reveal Kr-h1 and AstC-R are regulated by miR-277/34

The question then becomes, how significant is the finding of the revealed microRNA interaction, and could it really alter any biological activities or functions? Using the available homozygous loss of function of miR-277/34 cluster fly line (w[*]; Df(3R)mir-277-34-KO, TI{w[+mW.hs]=TI}mir-277-34-KO/TM3, P{w[+mC]=GAL4-twi.G}2.3, P{UAS-2xEGFP}AH2.3, Sb[1] Ser[1]; Chen et al 2014), we found that the heterozygous knockout individuals displayed almost no lethality from the mutation, while the survivorship for the double knockout flies fell significantly with 77% of the double knockout flies dying within five days after eclosion (Figure 2A).

**Figure 2.**
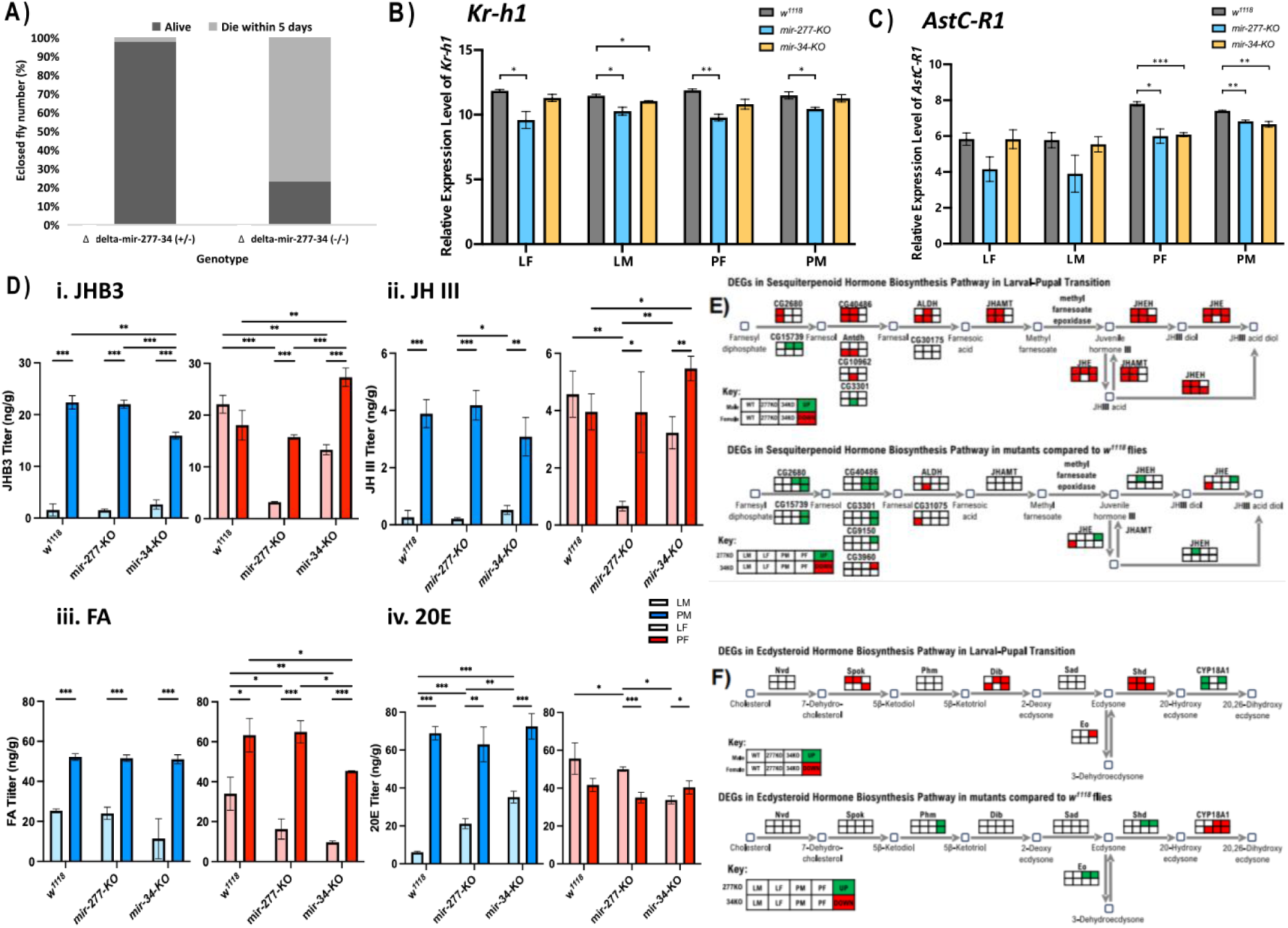
A) Mortality of heterozygous miR-277 and miR-34 double knockout and homozygous miR-277 and miR-34 double knockout flies after eclosion within 5 days; B) *Kr-h1* and C) *AstC-R1* relative expression level in larvae male (LM), larvae female (LF), pupae male (PM) and pupae female (PF) of *w^1118^*, *miR-277-KO*, and *miR-34-KO* flies; D) i. JHB3, ii. JH-III, iii. FA and iv. 20E hormone titre of *w^1118^*, *miR-277-KO* and *miR-34-KO* for LM, LF, PM and PF; E) Differentially expressed genes showing up-(green) or down-regulation (red) in the sesquiterpenoid hormone biosynthesis pathway in *w^1118^*, *miR-277-KO*, and *miR-34-KO* flies; G) Differentially expressed genes showing up-(green) or down-regulation (red) in the ecdysteroid hormone biosynthesis pathway in *w^1118^*, *miR-277-KO*, and *miR-34-KO* flies.

To fully understand how microRNA cluster miR-277/34 could be involved in the regulation of sesquiterpenoid via non-biosynthetic pathway genes in the fly, we utilised CRISPR/Cas9 to generate two individual knockout mutants for each of the microRNAs (i.e. miR-277 and miR-34). The mutant lines, which we named miR-277-knockout (miR-277-KO) and miR-34-knockout (miR-34-KO), were successfully created with confirmation by both Sanger sequencing and next-generation genome re-sequencing (Supplementary Figure S3A-B).

We then compared the expression of *Kr-h1* and *AstC-R1* of miR-277-KO, miR-33-KO, and between KO mutants and wild type flies using real-time PCR. The expression level of *Kr-h1* was found to be significantly downregulated in both sexes of larva and pupa miR-277-KO mutants; whereas a reduction was only noted in the male larvae for miR-34-KO mutants (Figure 2B).

On the other hand, expression level of *AstC-R1* was found to be upregulated during metamorphosis in wild-type (*w1118*) flies, and such change becomes insignificant in both miR-277-KO and miR-34-KO mutants (Figure 2C; Supplementary Figure 6). In addition, we found that the expression levels of *AstC-R1* in pupae of both miR-277-KO and mir-34-KO flies were significantly different to that of the *w1118* flies (Figure 2C). All these data demonstrated that *Kr-h1* and *AstC-R* are stage-specifically regulated by miR-277/34 *in vivo*.

### Sesquiterpenoid (JHB3, JHIII and FA) and ecdysteroid titres are altered differently between sexes in miR-277 and miR-34 knockout mutants

During larva-pupal transitions in flies, either the absence or excess of sesquiterpenoids results in abnormal phenotypes, including pupa lethality (Riddiford and Ashburner 1991; Riddiford et al 2010; Wen et al 2015). To assess the roles of miR-277 and miR-34 in regulation of sesquiterpenoid levels and other key metamorphosis hormone ecdysteroids in flies, we measured the titres of sesquiterpenoid hormones (JHB3, JH III, and FA) and ecdysteroid 20-hydroxylecdysone (20E) on the whole body of late L3 larvae and early pupae of both males and females of miR-277-KO, miR-34-KO, and *w1118* (as control) (Supplementary Table S6.).

In both the sesquiterpenoid and ecdysteroid titres measurement, we observed differences between males and females in the microRNA knockout mutants as well as the controls (Figure 2D). For example, we could only observe significant differences of sesquiterpenoid level changes during metamorphosis in females but not males between the control, miR-277-KO, miR-34-KO (Figure 2D). These data showed that both sesquiterpenoids and ecdysteroids were altered by miR-277 and miR-34.

### Members of miR-277/34 regulate different sesquiterpenoid and related gene pathways

To characterize global gene expression changes triggered by losses of miR-277 and miR-34, we compared the transcriptomes of RNA extracted from wandering 3^rd^ instar larvae (L3) and white prepupae from both sexes of miR-277-KO and miR-34-KO (Supplementary Figure S8-10). In the gene pathway enrichment analyses, we revealed that the insect hormone biosynthetic pathways were enriched in both control and miR-277-KO males (Figure 2F, Supplementary Figure S8-9). In addition, in evaluation of gene expression of those involved in sesquiterpenoid and ecdysteroid hormones biosynthetic pathways during larvae-pupae transition, we also found that miR-277 and miR-34 affect their gene expression to different degrees and trends in agreement to the hormonal measurement (Figure 2F and 2G).

The locus of microRNA cluster miR-277/34 has a common promoter and produces three noncoding RNA transcripts including CR43459-RB and CR43459-RC. CR43459-RB will encode miR-277, while CR43459-RC will encode both miR-277 and miR-34 (Figure 3A). These two noncoding RNAs CR43459-RB and CR43459-RC have differ expression profile, with CR43459-RC has a specific expression only at the adult male, and CR43459-RB has higher expression in embryonic, early larval, pupae, and adult stages but drops to low levels only during the late pupal stage before metamorphosis (Figure 3A). These expression profiles suggesting that miR-277 and miR-34 maybe functioning in a cooperative manner.

**Figure 3.**
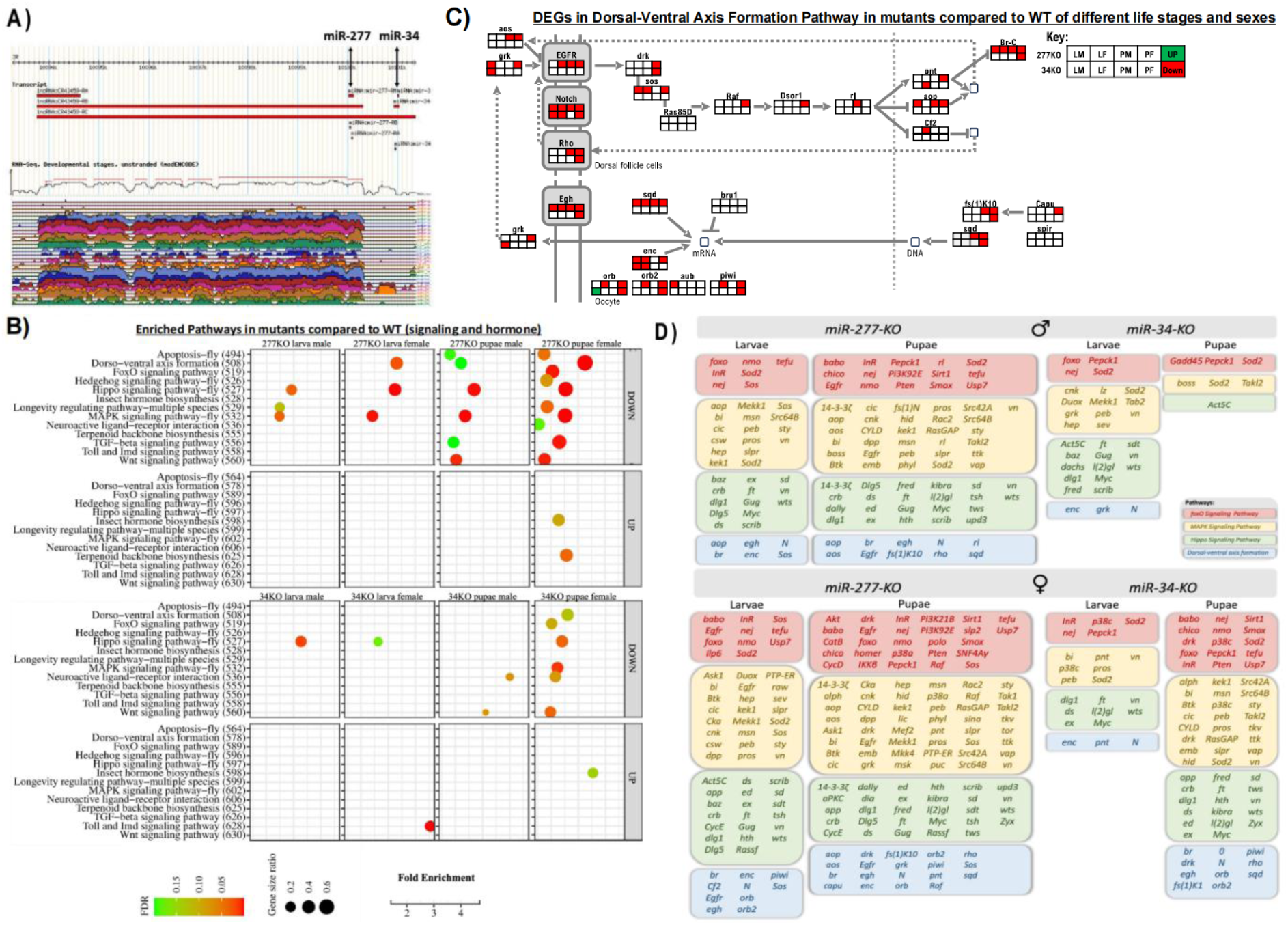
A) Expression level of transcripts in the miR-277/miR-34 locus retrieved from Flybase (version FB2023_0); B) Enriched gene pathways in *miR-277-KO* and *miR-34-KO* flies when comparing to *w^1118^* at different stages in male and female; C) Differentially expressed genes showing up-(green) or down-regulation (red) in *dorsal-ventral axis formation pathway* in *miR-277-KO* and *miR-34-KO* in males and females at larvae and pupae stages. C) Genes showing downregulation in the *FoxO signalling*, *MAPK signalling, Hippo signalling*, and *dorsal-ventral axis formation pathways* in *miR-277-KO* and *miR-34-KO* in males and females at larvae and pupae stages.

To understand which target genes can be regulated by members of this miR-277/34 cluster, we first compared the enriched gene pathways in the miR-277-KO and miR-34-KO mutants at different developmental stages and sexes when comparing to the wild-type *w1118* (Supplementary Figure S8-9). We found that there are both similar and unique gene pathways regulated by miR-277 and miR-34. For example, in the miR-277-KO mutants, alterations to the *Hippo signaling*, *MAPK signaling*, *TGF-beta signaling*, *FoxO signaling*, *Wnt signaling*, and *Dorso-ventral axis formation* gene pathways related to the sesquiterpenoid hormones are revealed (Figure 3B-D; Supplementary Figure S11-13); while the *Broad complex* (*Br-C*) which is a downstream transcription factor of the ecdysteroid hormonal pathway is significantly downregulated in miR-277-KO flies but only in miR-34-KO pupae females (Figure 3C), suggesting that the microRNA members in this miR-277/34 cluster regulate the sesquiterpenoid and related gene pathways in a cooperative manner (summarized in Figure 3B and D).

### Differential gene pathways regulated by miR-277/34 between sexes during metamorphosis

Further dissection of the above altered gene pathways in miR-277-KO mutants revealed that the *Hippo signaling* and *MAPK signaling pathways* are mutually downregulated regardless of sexes and life stages, whereas the *TGF-beta signaling* and *FoxO signaling* are downregulated exclusively in female pupae (Figure 3B, Supplementary Figure S8). In addition, *Wnt signaling pathway* is often downregulated in pupae, while the *Dorso-ventral axis formation* pathway is significantly downregulated in females but not in males. This data prompted us to consider that miR-277 while having conserved regulatory roles of sesquiterpenoid and related pathways, could also play divergent roles between sexes and metamorphosis stages.

Based on the new findings, we have compared differentially expressed genes during metamorphosis (i.e. larvae vs pupae) in different sexes between miR-277-KO, miR-34-KO, and controls. Genes in the *Toll and Imd signaling pathway*s were mainly downregulated in miR-277-KO females and miR-34-KO males, but slightly upregulated in miR-277-KO males through larvae-pupae transition (Figure 4A, Supplementary Figure S1 and S14). Interestingly, there are more male-specific downregulated gene pathways exclusively revealed in miR-277-KO males during larvae-pupae transition, including the *arginine and proline metabolism, fatty acid degradation, beta-alanine metabolism*, *cysteine and methionine metabolism*, *fatty acid degradation, pantothenate and CoA biosynthesis*, and *valine leucine and isoleucine degradation* (Figure 4A and B, Supplementary Figure 1). On the other hand, there are much fewer enriched downregulated gene pathways in miR-34-KO mutants during metamorphosis, such as the *other types of O-glycan biosynthesis* gene pathways which were specific to only the male flies (Supplementary Figure 1, Supplementary Table S7). These data added up that the members of microRNA cluster miR-277/34 also play differential gene regulatory roles between sexes during metamorphosis (Figure 4B).

**Figure 4.**
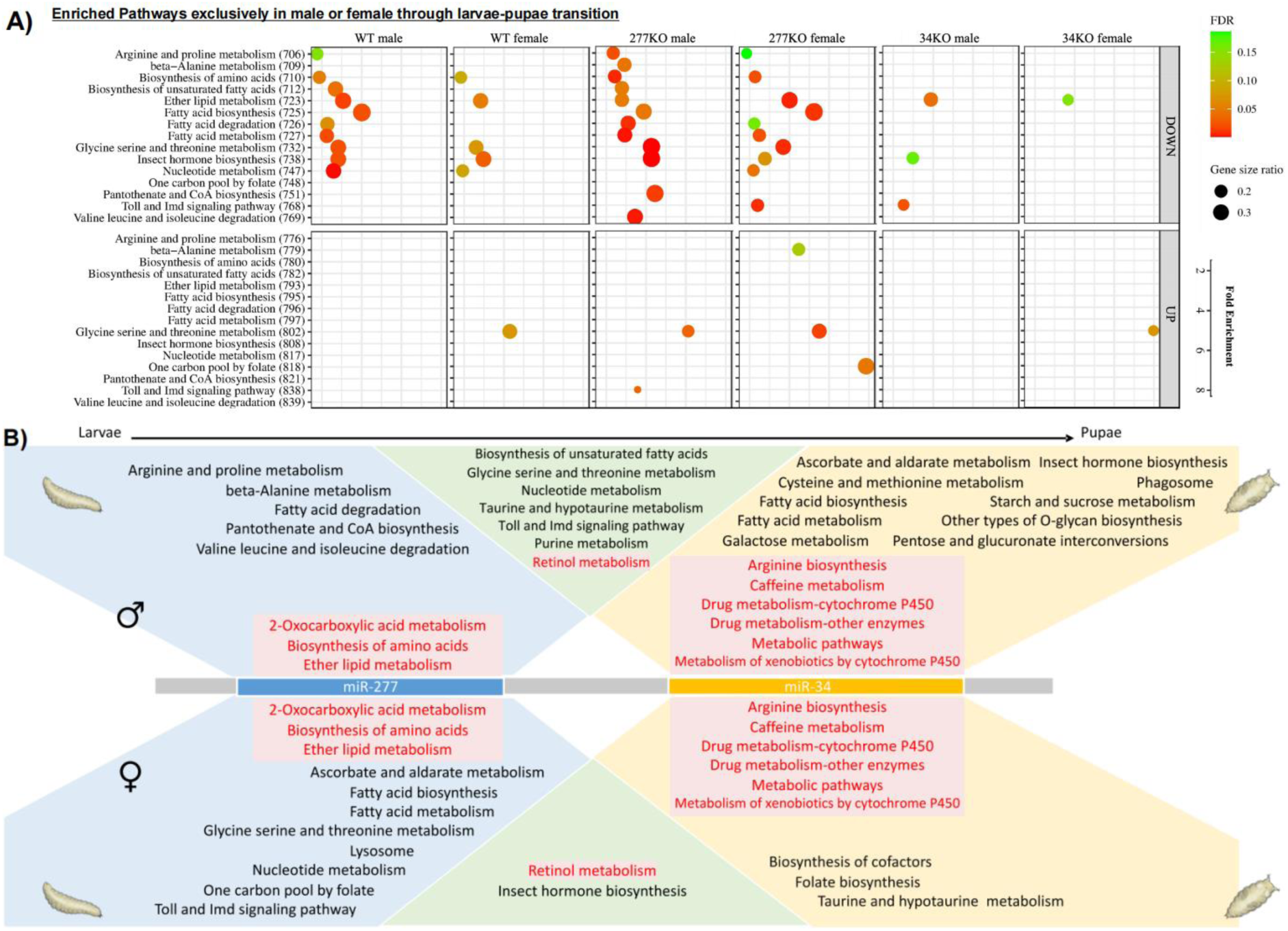
A) Enriched gene pathways in males and females of *miR-277-KO* and *miR-34-KO* that are unique to one sex during larvae-pupae transition; B) Schematic summary of gene pathways enriched with differentially expressed genes during larvae-pupae transition.

## Discussion

Animals display sexual dimorphism, and in the fruit fly *Drosophila*, sex-specific differences in body morphology, behaviour, and gene regulation have been previously described (e.g. Han et al 2022; Wang et al 2011; Coyne and Oyama 1995). After fly eclosion, different titres of ecdysteroids have been revealed in males and females potentially due to body size differences (Bownes et al 1984; Testa et al 2013). There is a plethora of studies showing sex-biased expression of micoRNAs across metazoans, however, there remains limited examples demonstrating how microRNAs could contribute the differences in insect sexes during metamorphosis and related hormonal control. We have first generated transcriptomes of both mRNA and microRNAs from both sexes in late larvae and early pupae stages, carried out genome-wide *in silico* prediction, *in vitro* luciferase assays, and *in vivo* pull-down assay, and narrowed down a microRNA candidate miR-277, as part of the miR-277/34 cluster, which can potentially involve in regulation of AstC-R1 and Kr-h1 transcripts. Previous studies have demonstrated miR-277 play various functional roles in flies, including circadian rhythm (Anna et al 2023), life span (Esslinger et al 2013), neurodegeneration and neuromuscular control (Tan et al 2012; Deshpande et al 2024; Cerro-Herreros et al 2016), homeostasis (Jones et al 2013; Li et al 2020), fatty acid metabolism (Zipper et al 2022), and amino acid metabolism (Esslinger et al 2013; Stark et al 2003). On the other hand, miR-34 has also been shown to play other roles in flies such as in autophagy, brain development, life span, survivorship, and regulation of ecdysone (Liu et al 2012; Lai et al 2016; Kennerdell et al 2018, McNeill et al 2020; Srinivasan et al 2022; Lai et al 2020; Xiong et al 2016). Together with the correlated expression data (AstC-R1, Kr-h1, miR-277 and miR-34), *in vivo* mutants construction, hormone measurement and transcriptomic analyses, this study strongly demonstrated that miR-277 and miR-34 can play cooperative biological roles in sesquiterpenoid hormones in the fly sexes during metamorphosis.

Unlike plants and fungi, metazoans including insects usually have significant proportion of microRNAs that are clustered in their genomes (e.g. Bartel 2018; Tanzer and Stadler 2004). Members of these microRNA clusters often transcribed as a single unit of polycistronic transcript regulated by a single promoter (Kozomara and Griffiths-Jones 2014). For instance, misregulation of clustered microRNAs could lead to disease and cancer formation in humans (Dambal et al 2015; Ventura et al 2008; Kim et al 2009), and thus microRNAs in the same cluster are proposed to possess similar targeting properties or regulate genes in the same pathway (e.g. Hausser and Zavolan 2014). In *Drosophila*, microRNA cluster miR-6/5/4/286/3/309 has been demonstrated that new members can reinforce the gene regulatory network of existing microRNAs (Qu et al 2020), and let-7/miR-125 cluster are also well known to be controlled by ecdysteroid hormones (e.g. Garbuzov and Tatar 2010). This study now further provided evidence for functional constraints in microRNA clustering in insects, and in this case of miR-277/34, via regulating different hormonal and related gene pathways in different sexes during metamorphosis.

## Materials and Methods

### Animal husbandry, sex differentiation, and transcriptomes

D. melanogaster w1118 was reared at 25 C, 45-50% relative humidity and 14:10 hour light-dark cycle. Sexes of wandering L3 larvae instar were differentiated under the microscope. Total RNA was extracted from different sexes of L3 larvae instar and white pre-pupae stages with TRIzol® Reagent (Thermo Fisher Scientific, US). The integrity and quantity of RNA was verified through gel electrophoresis and NanoDrop One/OneC spectrophotometer (Thermo Fisher Scientific, US). Total RNA was submitted to Novogene Hong Kong for library construction and sequencing on the Illumina Novaseq6000 (PE150) platform. Raw reads were trimmed and filtered using Trimmomatic (version 0.39, with parameters ’ILLUMINACLIP: TruSeq3-PE.fa:2:30:10 SLIDING WINDOW:4:5 LEADING:5 TRAILING:5 MINLEN:25’) (Bolger et al 2014), while contamination were removed using Kraken2 (version 2.0.8) (Wood et al 2019). The processed reads were then aligned using HISAT2 (v2.0.5) (Kim et al 2019) and gene matrix count tables were generated using StringTie (v2.1.1) (Kovaka et al 2019) with references downloaded from Flybase (version FB2023_02). Details can be found in Supplementary Table S8.

In addition, small RNA was also extracted from different sexes of L3 larvae instar and white pre-pupae stages with mirVana™ miRNA Isolation Kit (Ambion). The integrity and quantity of RNA was checked using an Agilent 2100 Bioanalyser (Agilent RNA 6000 Nano Kit), and submitted to BGI Hong Kong for HiSeq small RNA library construction and 50-bp single end sequencing. Small RNA sequencing reads were analysed as previously described (Qu et al 2020). In brief, adapters were trimmed using cutadapt (version 1.11) (Martin 2011) and quality control was performed using fastqc (Andrews 2010). Qualified reads within 18 and 26 bp were used to generate a matrix count table using the ’quantifier.pl’ module of the mirDeep2 package (Friedländer et al., 2012), with ’-g 0’ to allow for 0 mismatch when mapping reads to precursors downloaded from miRBase (version 22.1) (Kozomara et al 2019). The resulting count matrix was used for expression analyses in edgeR (Robinson et al 2010). Clean reads were also aligned to precursor sequences using Bowtie (Langmead et al 2009) with the parameters "-l 18 -v 0 -a --best --norc --strata" followed by manual inspection and checking. Details can be found in Supplementary Table S8.

### MicroRNA target prediction and validation (dual-luciferase reporter, LAMP, and quantitative microRNA Taqman assays)

MicroRNA target prediction and in vitro validation were carried out as previously described (Qu et al 2017, 2020). In brief, mature microRNA sequences were retrieved from miRbase (version 22.1) (Kozomara et al 2019), and used to predict targets using the miRanda algorithm (Enright et al 2003). In addition, target prediction was also performed with TargetScanFly (release 7.2) (Agarwal et al 2018).

To validate the predicted microRNA targets, a dual-luciferase reporter assays were used (Qu et al 2017, 2020). MicroRNAs were amplified from D. melanogaster with amplicons cloned into pAC5.1 vector (Invitrogen), while gene targets were cloned into psi-check2 vector (Promega). All constructs were sequenced to confirm their identities. Drosophila S2 cells were kept at 23-25℃ in Schneider Drosophila medium with 10% (v/v) heat-inactivated fetal bovine serum and 1:100 penicillin–streptomycin (Gibco, Life Technologies). The pAC5.1-microRNA (200 ng) and psi-check2 vector (100ng) were transfected into Drosophila S2 cells using Effectene (Qiagen) per the manufactures’ instructions. Luciferase activities were measured at 36-48 hours post-transfection using Dual-Luciferase Reporter Assay System (Promega) using Tecan Spark Multimode Microplate Reader (Eastwin International Trading Ltd).

Further validations were also carried out for miR-277 and miR-34 with labelled microRNA pull down assays (LAMP) as well as expression level analyses (TaqMan™ MicroRNA Reverse Transcription Kit and TaqMan™ MicroRNA Assay (Applied Biosystems), Assay IDs are 464452_mat, 463733_mat, 000298, and 463695_mat) as previously described (Qu et al 2017, 2020). Biotin-labelled oligos synthesized by IDT company while the Dynabeads M-280 streptavidin were obtained from the Invitrogen. The PCR was performed with the StepOnePlus™ Real-Time PCR System (Applied Biosystems). Sequences of oligos are provided in Supplementary Table S9.

### miR-277 and miR-34 knockout mutants generation and confirmation

To prepare the CRISPR deletion constructs of miR-277 and miR-34, the corresponding target sequences and sequences for homologous recombination repairing were cloned into the pU6-3-Bsal-gRNA and pHD-DsRed-attP vectors (primer information is provided in Supplementary Table S9). Constructs were sequenced prior to injection into D. melanogaster w1118 embryos. D. melanogaster homozygous miR-277 and miR-34 CRISPR knockout mutants were screened and selected based on markers and phenotypes. Genomic DNA of each mutant line was extracted with PureLink DNA extraction kit (Invitrogen) following the manufacturer’s protocol. PCR was performed with the primer sequences provided in the supplementary Table S9 to confirm that the respective microRNA has been taken out. In addition, whole genome resequencing was also performed on the these mutants on Illumina HiSeq platform at the Novogene Hong Kong. Sequenced raw reads were trimmed and cleaned using Trimmomatic (v0.39) (Bolger et al., 2014) and Kraken 2 (Wood et al., 2019) and aligned to the D. melanogaster reference genome downloaded from Flybase (r6.45) using BWA-MEM (Li, 2013) with parameters "-M -R", and the alignment was checked manually using samtools tview.

### Ecdysteroid hormone measurement

Total body ecdysteroids were extracted from w1118, mir-277 knockout mutants (miR-277-KO) and mir-34 knockout mutants (miR-34-KO) L3 larvae and early pupae from both sexes (n=3, 50 individuals per sample using methanol (Furuta et al., 2010), and the levels were determined using the 20-hydroxyecdysone (20E) assay described previously (Kai et al., 2021).

### Sesquiterpenoid hormone measurement

Total body sesquiterpenoid juvenile hormones were collected from w1118, miR-277-KO and miR-34-KO L3 larvae and early pupae from both sexes (n=3, 50 individuals per sample) and freeze-dried. The titre of FA, JH III and JHB3 were determined by adapting a protocol from a previous study (Kai et al. 2018). In brief, the freeze-dried homogenates were transferred to 10 ml glass centrifuge tubes containing 1 mL acetonitrile, 1 mL 0.9% (w/v) sodium chloride solution and 10 ng of JH III-D3 as an internal standard, ultrasonicated for 1 min, and then vortexed and extracted twice with 2 mL hexane. The hexane phase (upper layer) was removed and transferred to a new glass vial, and then dried under nitrogen flow. The residue was dissolved in 1 mL acetonitrile. The measurement of FA, JH III and JHB3 was determined based on the previous method with little modifications reported by Ramirez et al. (2020). The assay was performed on the Acquity UPLC (Waters) system connected to the AB SCIEX 5500 triple-quadrupole mass spectrometer (Framingham, MA, USA). Isolated the target analytes on a BEH C18 column (100 mm* ×2.1 mm, 1.7 µm particle size) maintained at 30°C. The mobile phases were methanol (A) and 0.1% formic acid in ultrapure water (B), and the flow rate was 0.3 mL min−1. The gradient program started at 10% A hold on for 1 min, changed to 90% A (in 1–7 min), and decreased to 10% A (7–8 min). The injection volume was 8 µL. The ion spray voltage was set at 5500 V, 99.99% N2 was used as the desolvation/nebulizer gas, and 99.99% Ar as the collision gas. The temperature of the block source was maintained at 500°C, while the pressure of nebulizer gas and turbo gas was set at 50 psi. The curtain gas (CUR) pressure was 38 psi, and the collision gas (CAD) value was set to 8. AB Sciex v1.6 analyst software was used for controlling instruments, data acquisition and processing. The optimal MRM transitions and other parameters of UPLC-MS/MS for the test JHs were given in Supplementary Table S6.

### Transcriptomic analyses of mutants metamorphosis and between sexes

Total RNA was extracted from L3 larvae and pre-pupae of miR-277-KO and miR-34-KO flies using TRIzol® Reagent (Thermo Fisher Scientific) and sequenced on Illumina HiSeq platform. Raw reads were processed as above, and the processed reads were aligned using HISAT2 (v2.0.5) (Kim et al 2019). Gene matrix count tables were generated using StringTie (v2.1.1) (Kovaka et al 2019) with references downloaded from Flybase (version FB2023_02). Differentially expressed genes (DEGs) were identified using iDEP 1.0 (Ge et al 2018) to generate the hierarchical clustering heatmap, principal component analysis (PCA), enrichment analysis and enriched pathway analysis with default parameters. The DEGs were also analysed with ShinyGO (v0.77) (Ge et al 2020) for gene enrichment analyses. Details can be found in Supplementary Table S7.

## Availability of data and materials

The raw reads generated in this study have been deposited to the NCBI database under the BioProject accessions: PRJNA974543.

## Acknowledgments

This work was supported by the Hong Kong Research Grant Council General Research Fund (14100420). KKC was supported by a PhD studentship supported by The Chinese University of Hong Kong.

