## Supplementary Figure for "Insect metamorphosis is regulated differently between sexes by members of a microRNA cluster"

Supplementary Figure 1. Metamorphosis pupae vs larva

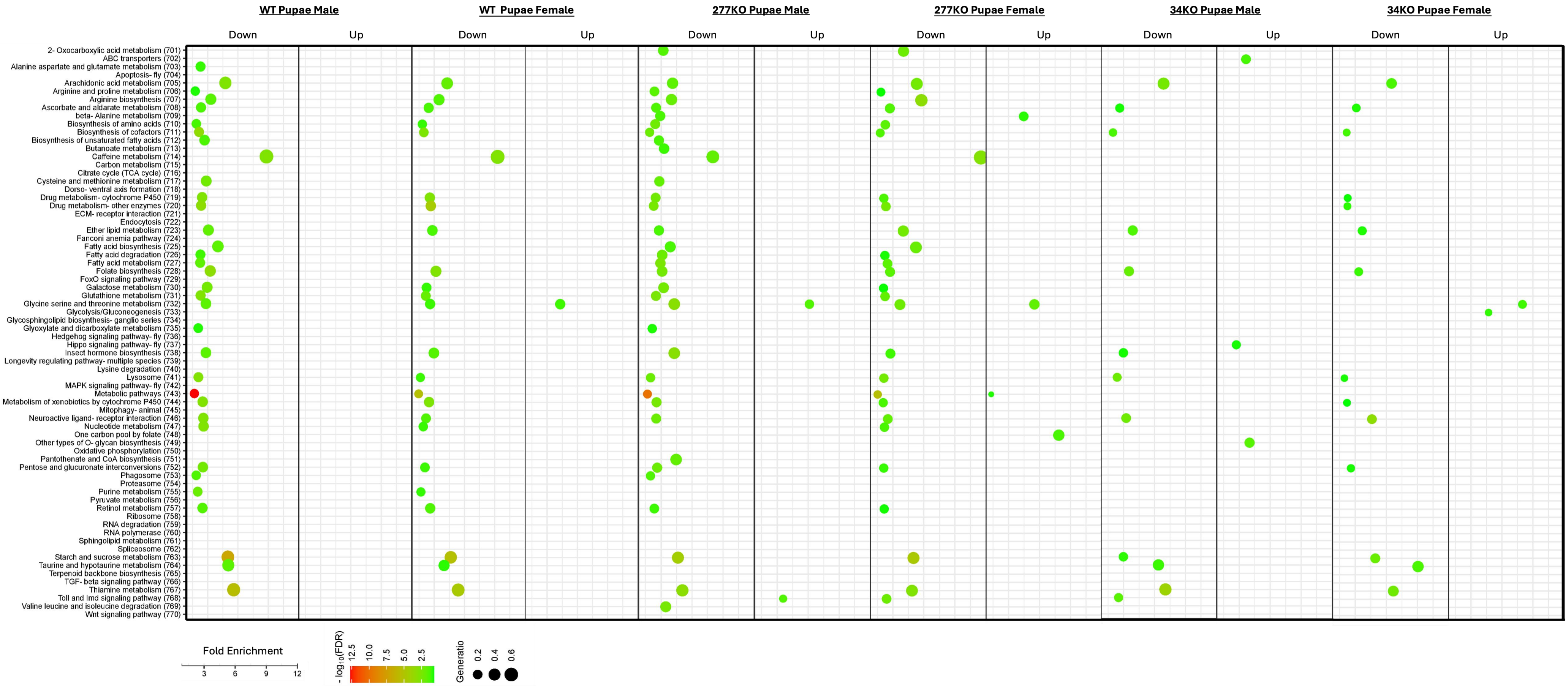

Supplementary Figure 2. Principal component analysis (PCA) plot of miRNA samples

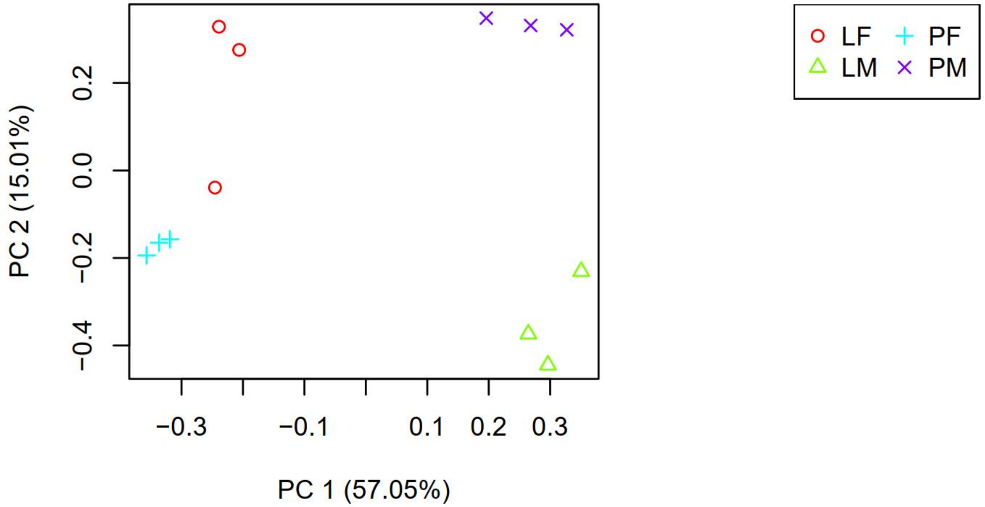


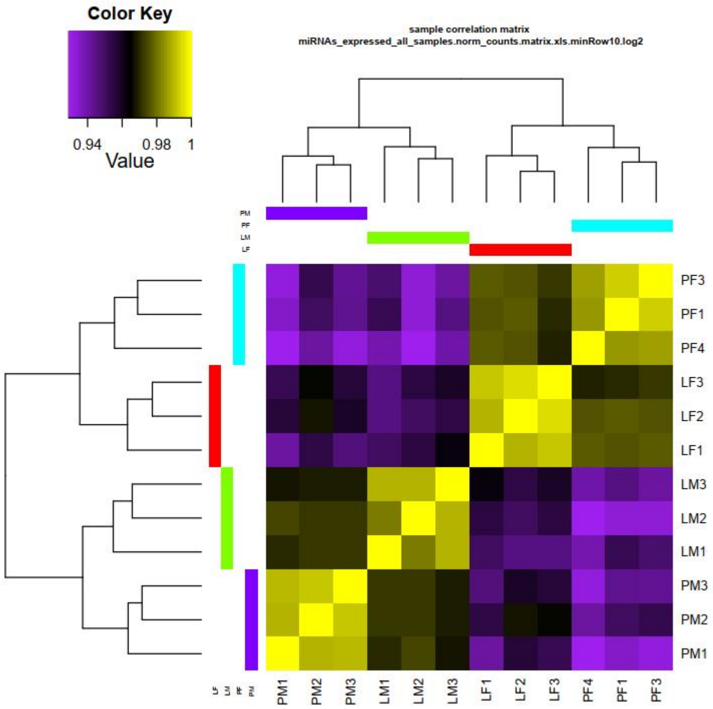

Supplementary Figure 3. Female vs Male of the miRNA expression

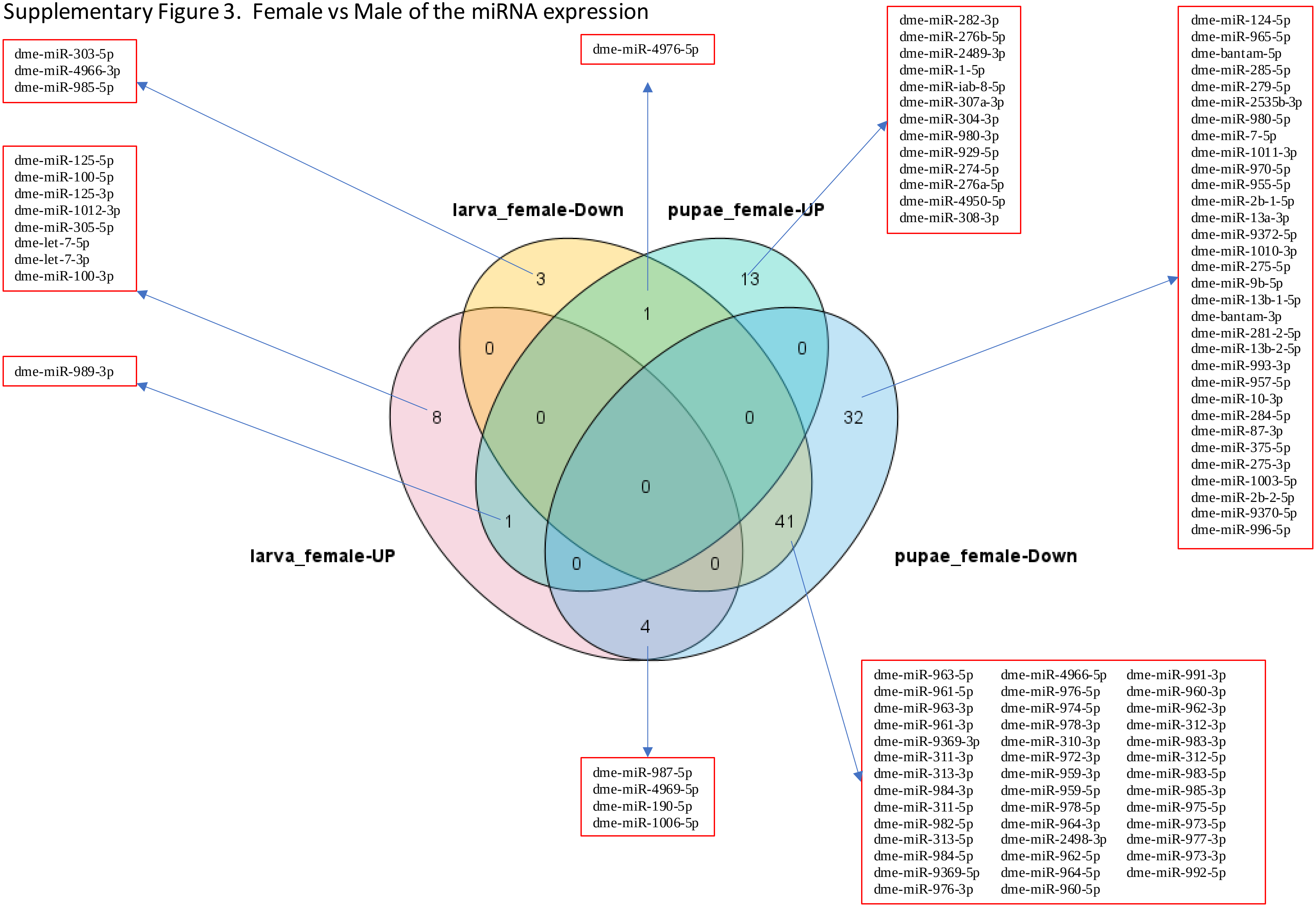

Supplementary Figure 4. pupae vs larva of the miRNA expression

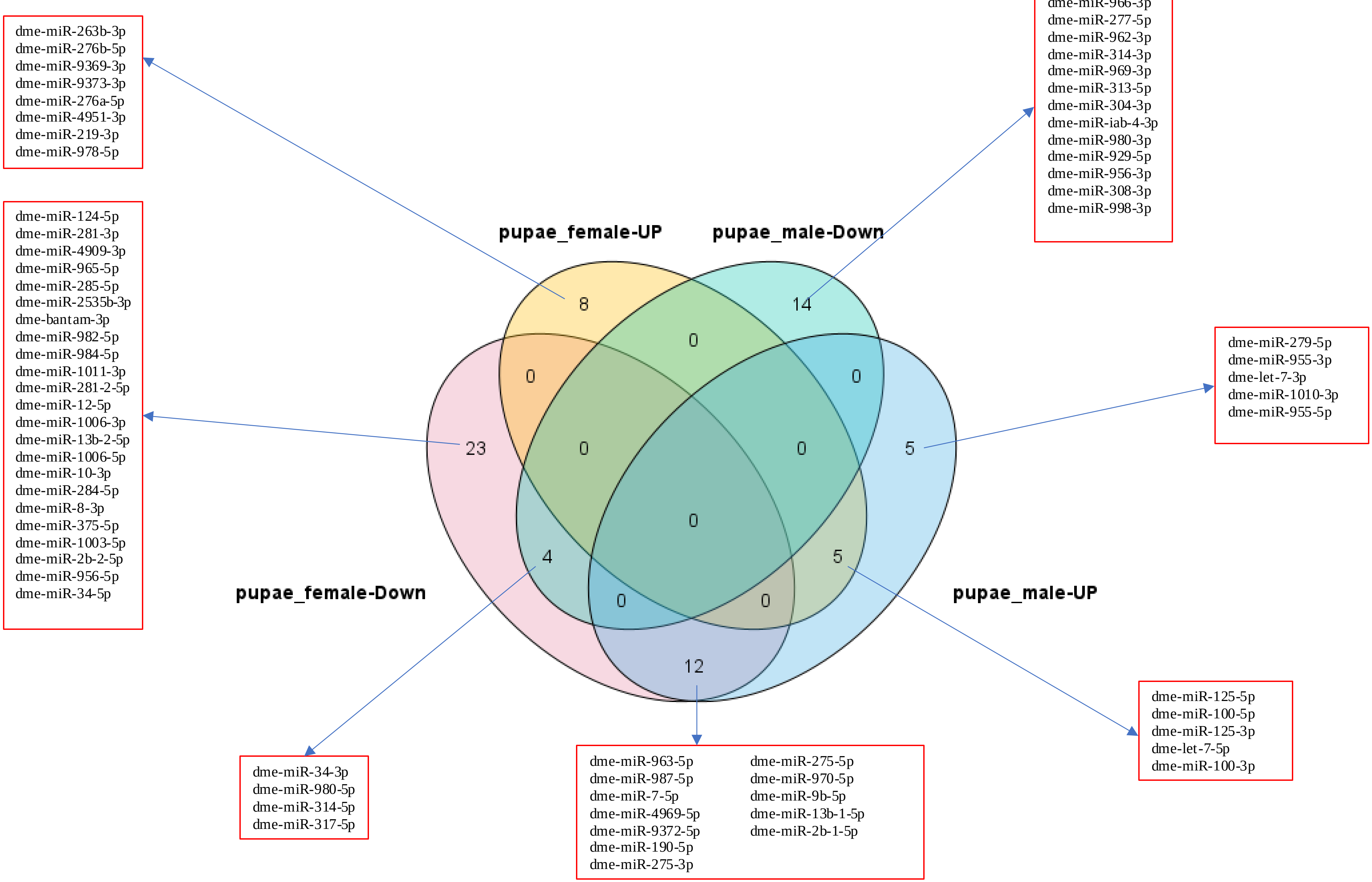

Supplementary Figure 5. *in silico* prediction on potential miRNA targets of the sesquiterpenoid system gene transcripts.

| Target Gene | JHDK | JHE | JHEDUP | JHBP | Chd64 | Kr-h1 | tai | AstC-R1 | AstC | AstC-R2 |
|---|---|---|---|---|---|---|---|---|---|---|
| Gene: miRNA *in silico* screening | miR-1000 | miR-1002 | miR-968 | miR-968 | miR-1000 | miR-1002 | miR-137 | miR-137 | miR-2279 | miR-283 |
| | miR-2283 | | | miR-991 | miR-1002 | miR-2279 | miR-2279 | miR-2279 | miR-2280 | miR-958 |
| | miR-315 | | | | miR-2279 | miR-263b | miR-2283 | miR-2280 | miR-263b | miR-985 |
| | miR-958 | | | | miR-2283 | miR-277 | miR-283 | miR-2283 | miR-927 | |
| | | | | | miR-315 | miR-315 | miR-985 | miR-277 | | |
| | | | | | miR-968 | miR-8 | | miR-8 | | |
| | | | | | miR-991 | miR-927 | | | | |
| | | | | | miR-985 | miR-991 | | | | |

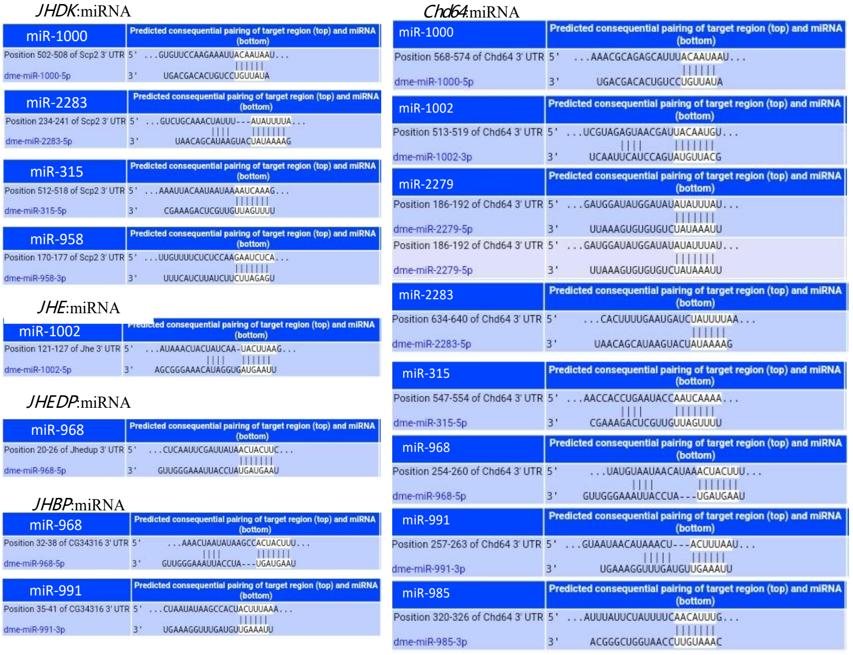


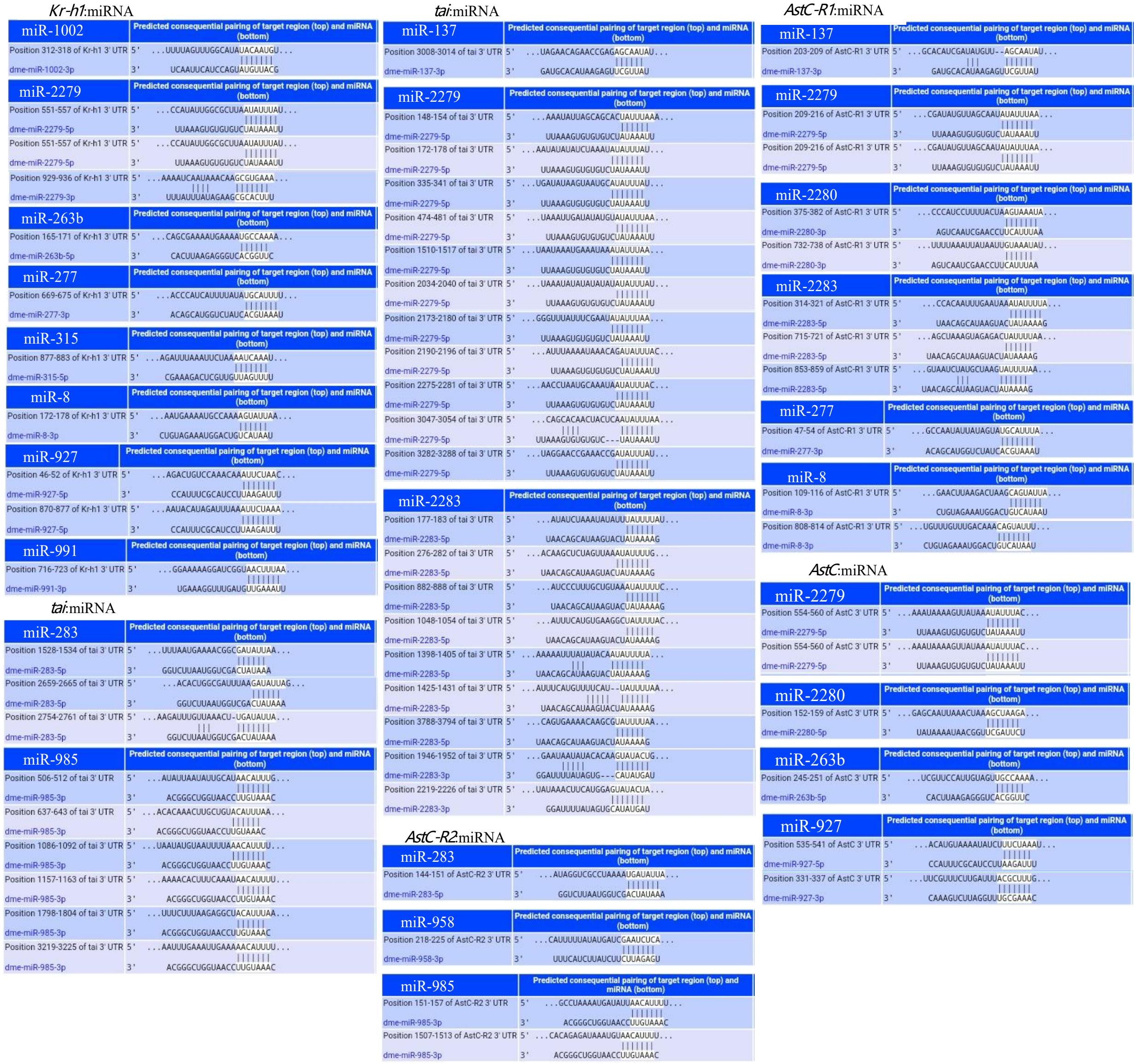

Supplementary Figure 6. Sanger sequencing of mutants and genomic position alignment of mir-277 and mir-34 from samples S277KO_6, S277KO_1 S34KO_9, and S8_3_1.

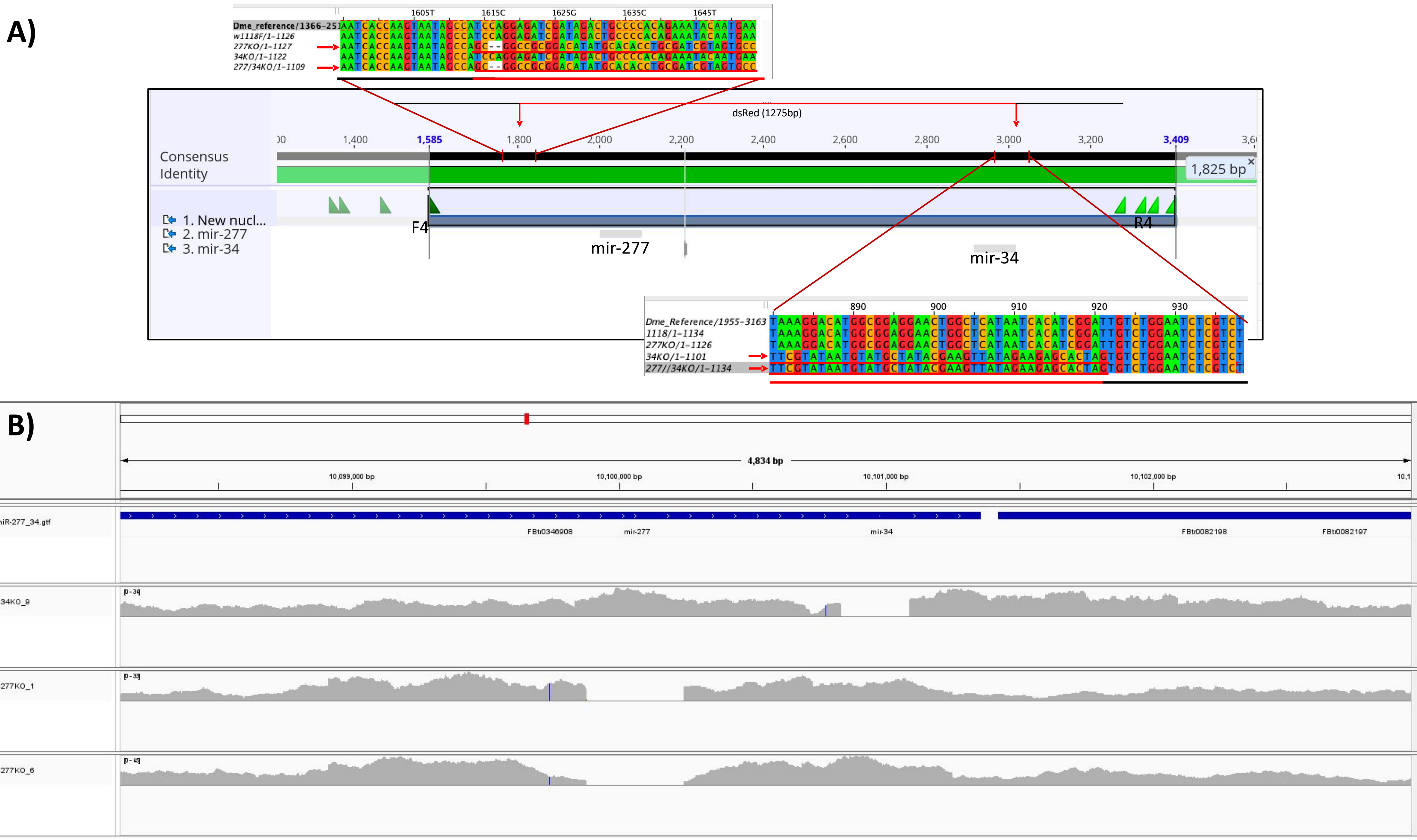

Supplementary Figure 7. The expression of *AstC-R1* in different sexes and stages, clustering according to the 3 genetic conditions ($w^{1118}$, miR-277-KO and miR-34-KO mutant).

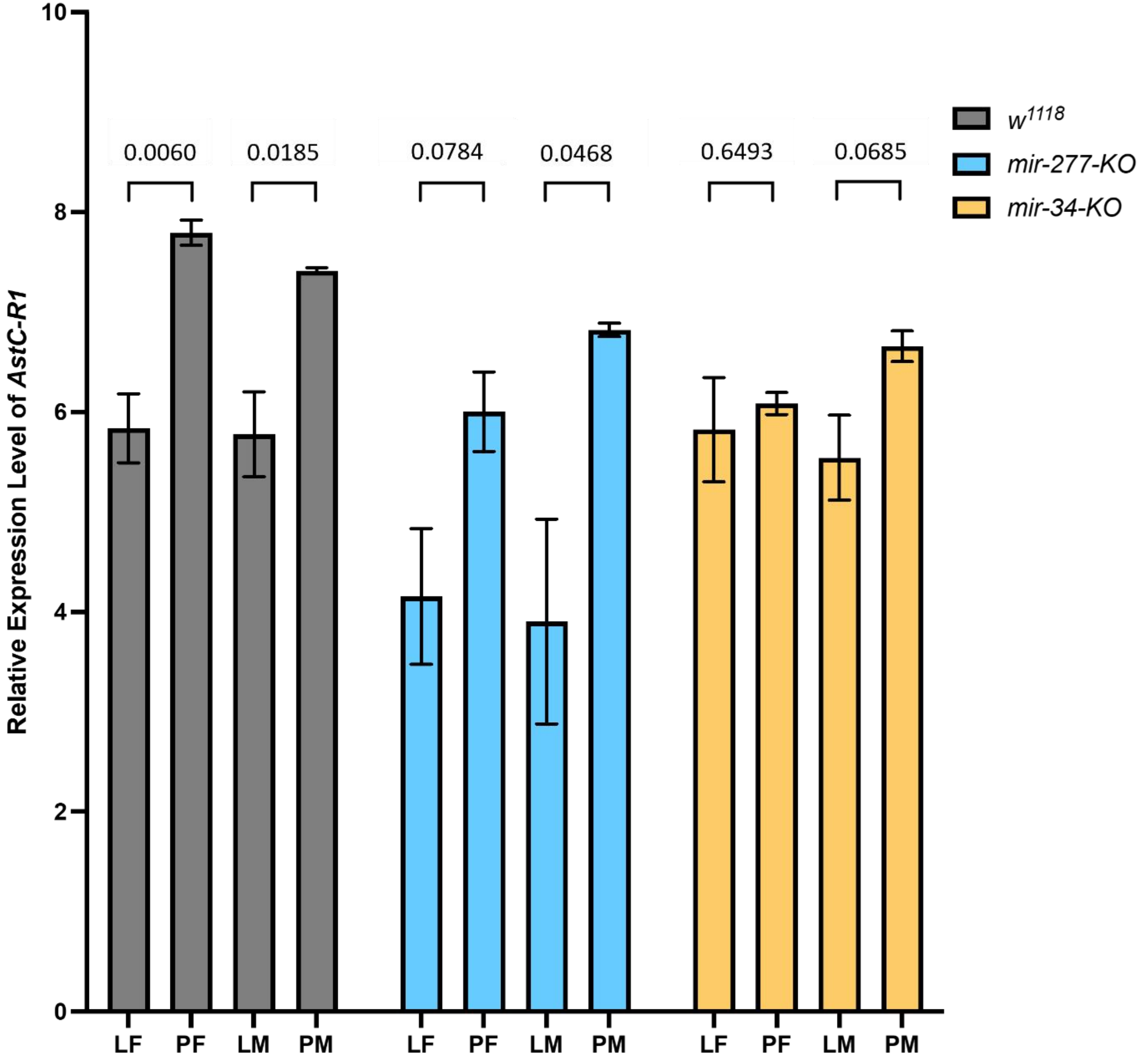

Supplementary Figure 8. Enriched gene pathways in miR-277-KO flies compared WT flies in different life stages and sexes

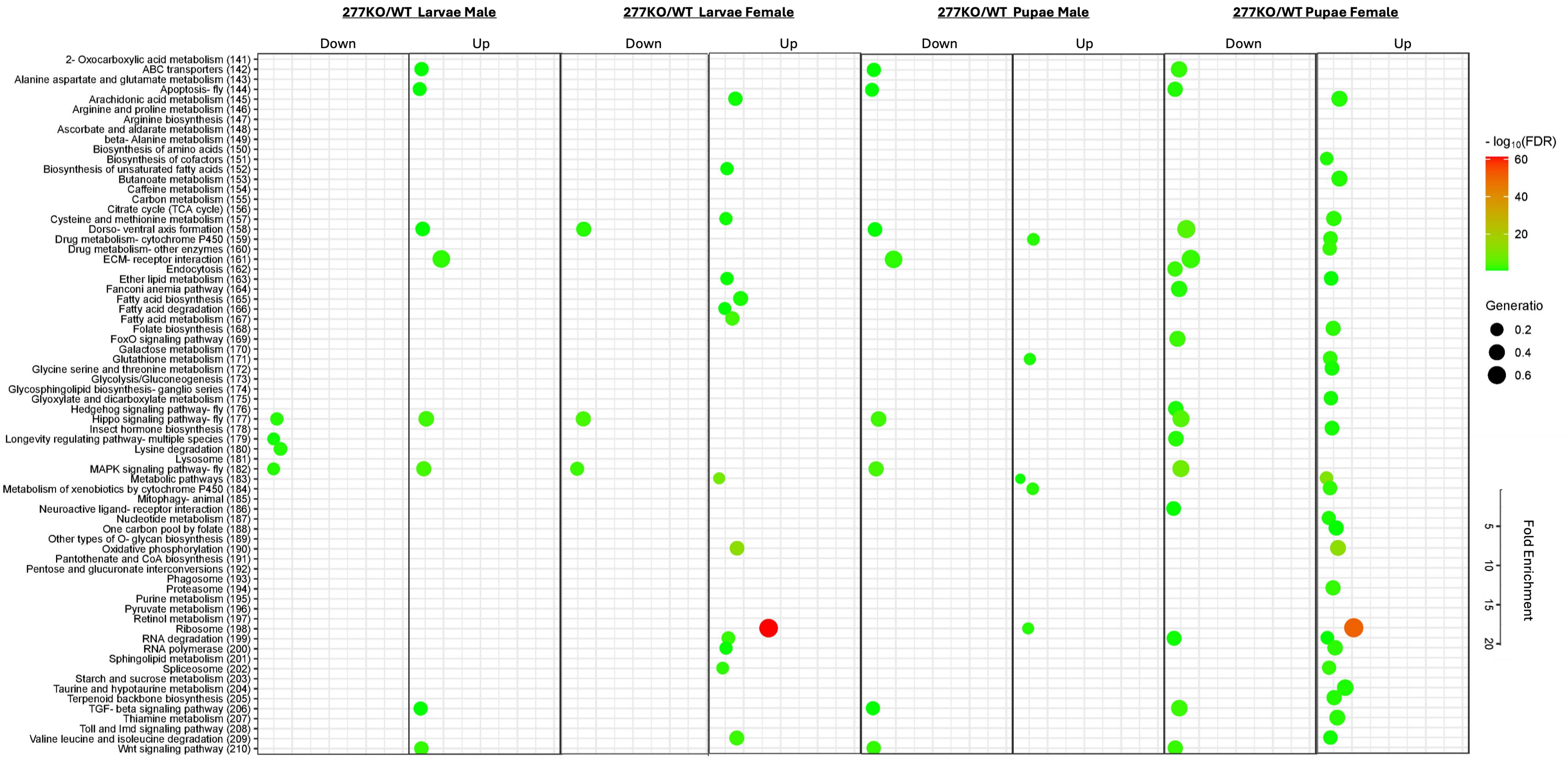

Supplementary Figure 9. Enriched gene pathways in miR-34-KO flies compared WT flies in different life stages and sexes

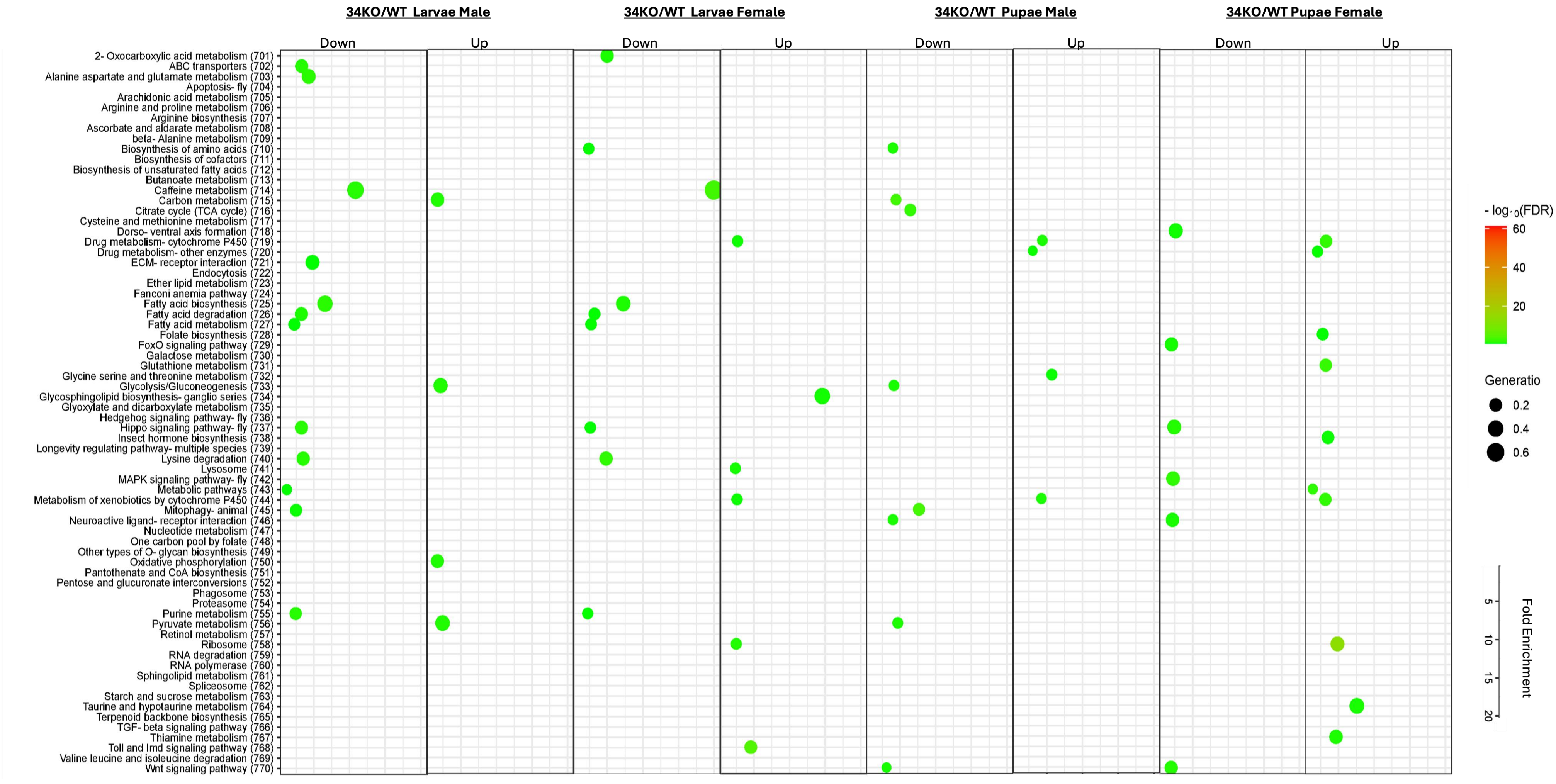

Supplementary Figure 10. Enriched gene pathways in female flies compared male flies in the WT and mutant flies at different life stages.

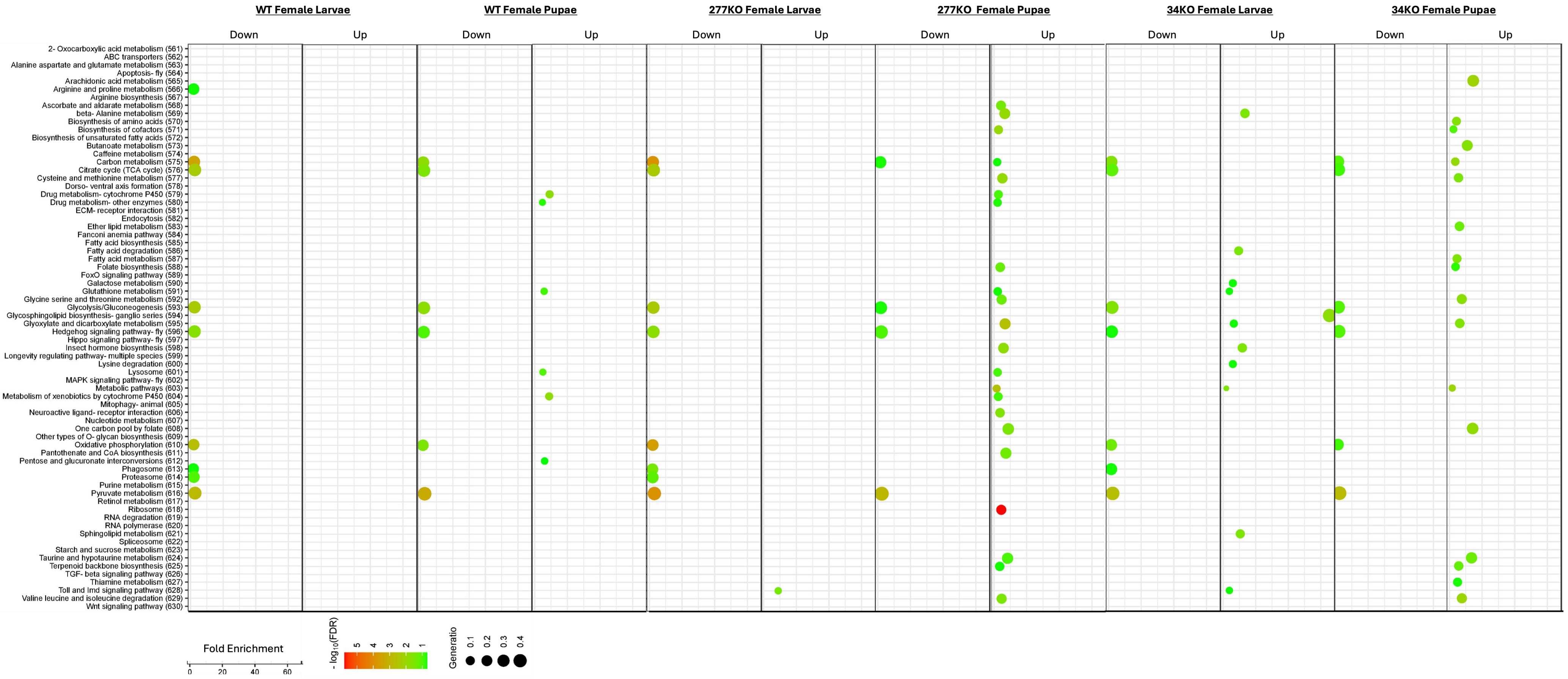

Supplementary Figure 11. DEGs enriched in mutants when compared with the WT flies in the MAPK signaling pathway.

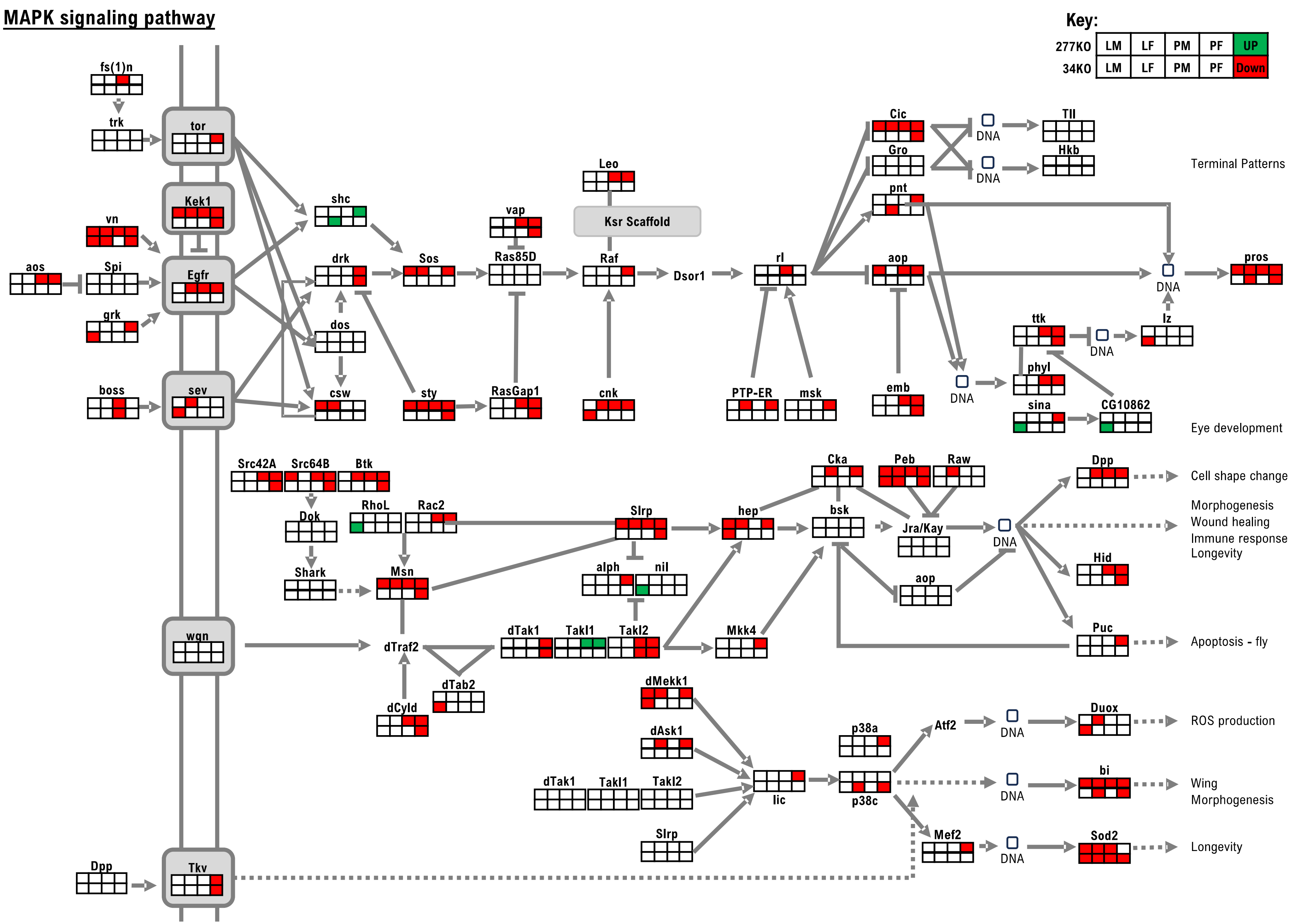

Supplementary Figure 12. DEGs enriched in mutants when compared with the WT flies in the Hippo signaling pathway.

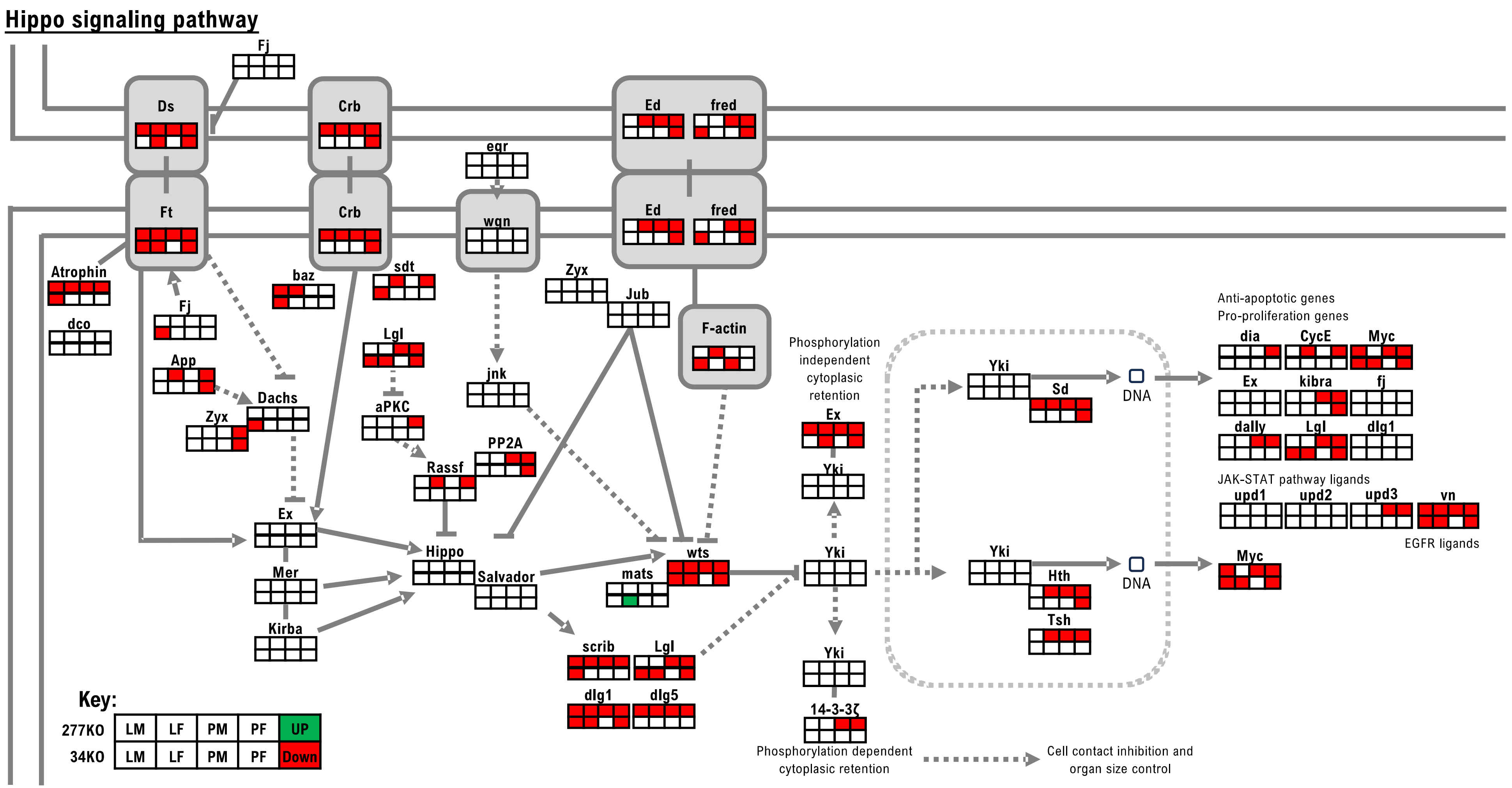

Supplementary Figure 13. DEGs enriched in mutants when compared with the WT flies in the FoxO signalling pathway.

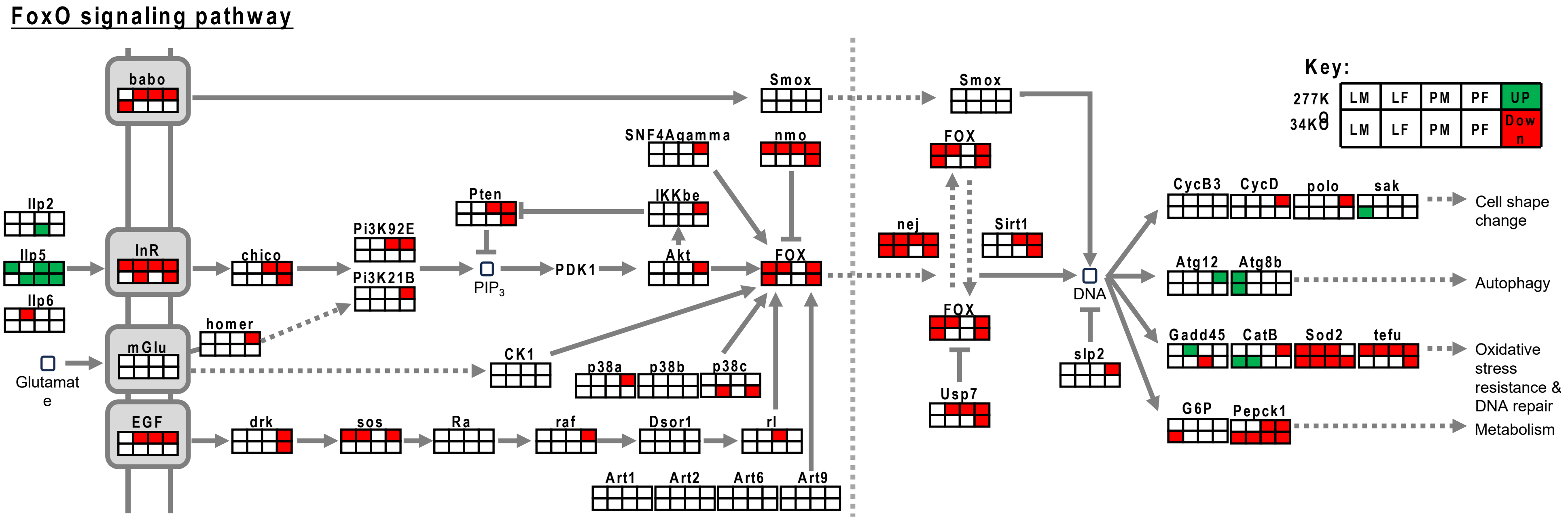

Supplementary Figure 14. DEGs enriched in the Toll and Imd Signaling Pathways through male and female larvae-pupae transition

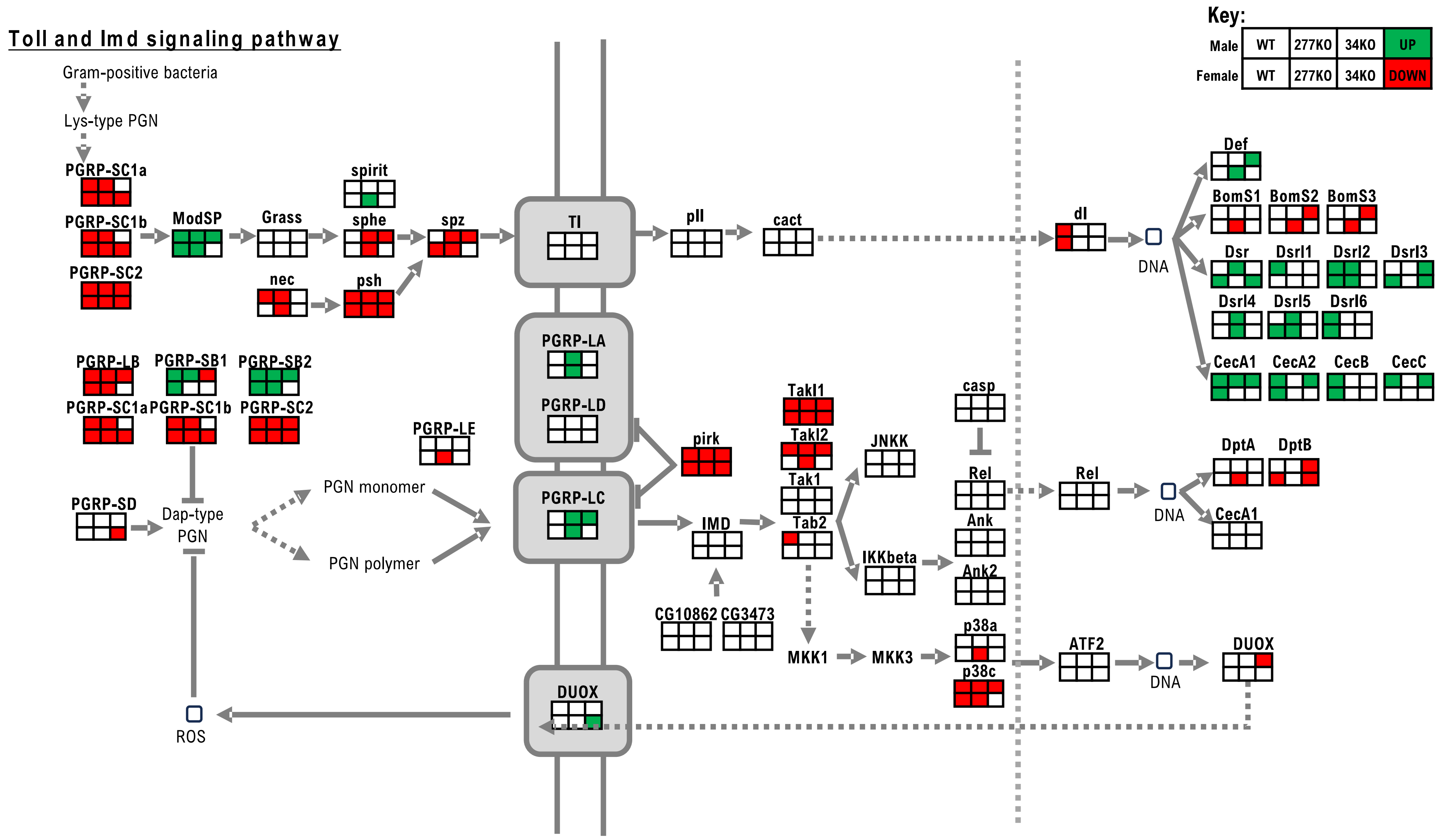
